# Study of the seasonal life cycle of *Lutzomyia longiflocosa* (Diptera: Psychodidae) under semi-controlled conditions in the department of Huila, Colombia

**DOI:** 10.1101/2023.07.11.548498

**Authors:** Jennifer Alejandra Vargas Durango, Gelys Igreth Mestre Carillo, Erika Santamaria Herreño, Miguel Angel Beltran Ruiz, Angela Cristina Zapata Lesmes, Astrid Geovanna Muñoz Ortiz, Jesús Eduardo Escobar Castro

## Abstract

**Background.** *Lutzomyia longiflocosa* is an insect vector associated with the transmission of Cutaneous Leishmaniasis. To establish the parameters of the life cycle of *L. longiflocosa* in semi-controlled conditions in a rural area of the Campoalegre municipality, Huila, Colombia. **Methodology/Principal Findings.** The life cycle of individuals of *Lutzomyia longiflocosa*, obtained from two cohorts of collected, fed, and individualized females, was monitored during two different times of the year (between February and August 2020 and between July 2020 and January 2021, respectively). Determining parameters associated with the fertility and fecundity, time and attributes of development and survival, and its association with abiotic variables. The average duration of Cycle 1 (C1) and Cycle 2 (C2) was 134.9 and 148.78 days, respectively. The gonotrophic cycle of parental females presented significant differences (p-value <0.05) between C1 and C2 (8.47 and 11.42 days) as well as between fecundity and fertility parameters. The number of days it takes the development of the immature stages between the two cycles studied, also showed significant differences in the larvae II (15.21 and 22.23), larvae III (11.93 and 17.56), and pupae (24.48 and 22.9) stages. During C1, the survival rate was higher and consistent with the productivity figures of adult individuals (F1), compared to C2. Fecundity and fertility values were significantly higher in C2. Finally, a significant correlation between the number of individuals and temperature was evidenced in C1 while, for C2, there was a negative correlation with precipitation. **Conclusions/Significance.** Significant differences were found in several biological and reproductive parameters between the two cycles monitored. The parameters of the life cycle of *L. longiflocosa* in its natural habitat would be influenced by environmental factors related to the annual seasonality in the sub-Andean rural area, conditioning the temporal distribution of this species and, consequently, the possible transmission of causative agents of cutaneous leishmaniasis.

**Author Summary:** Cutaneous leishmaniasis (CL) is a public health disease that affects more than 300 million people in tropical areas in the world and its causative agent is a parasite of the genus *Leishmania*. In recent decades, the cases in Latin America have increased, corresponding to 6% of cases in the tropical regions of the planet. An insect vector associated with the transmission of CL in these tropical and endemic areas is *Lutzomyia longiflocosa*. Therefore, it is very important to investigate the environmental parameters that can influence its life cycle and its transmission capacity and competition, aspects that are not fully known.

In Colombia, an endemic focus is located in the department of Huila, in a rural area of the municipality of Campoalegre, where this study was carried out. The authors found the relationship between some environmental parameters with the life cycle of *L. longiflocosa* under semi-controlled conditions in forested habitats, aspects that were not known until now. The impact of temperature, relative humidity, and precipitation on the gonotrophic cycle, fecundity, fertility, time of development and survival, was evidenced. This research provides information on the environmental conditions that directly affect the possible transmission of the agents that cause cutaneous leishmaniasis.

## Introduction

Cutaneous leishmaniasis (CL) is a zoonotic disease that has had an increase in cases in Latin America in recent decades, becoming a public health problem [1]. In 2020, Colombia reported the second highest number of cases in Latin America with a total of 6,161 and an incidence rate of 23.34 cases per 100,000 inhabitants [2].

CL vectors are represented in Colombia by 135 species of the genus *Lutzomyia* with a wide distribution in most geographical areas of the country, of which seven have been found naturally infected with *Leishmania spp* parasites [3,4,5,6]. Insects of the genus *Lutzomyia*, specifically, of the verrucarum group, townsendi series (*L*. *longiflocosa*, *L. spinicrassa, L. torvida, L. tonwsendi, L. quiasitonwsendi*, *and L. youngi*) have been frequently associated with cases of LC in mountainous areas of Colombia, Venezuela, and Costa Rica, mainly at elevations above 1,500 meters above sea level (m.a.s.l.), and in areas where there are ecological disturbances associated with the expansion of monocultures such as coffee [7,8,9].

Between 1990 and 2014, the highest number of cases of LC was reported in the sub-Andean geographical area of the Magdalena River Valley, amid 1,000 and 2,000 m.a.s.l., mainly in the departments of Huila and Tolima, where the *Lutzomyia longiflocosa* species [10] was considered the most likely vector in this area [11,12]. In these departments, *L. longiflocosa* is dominant with abundances >90% in the municipalities of Neiva, Baraya, Rivera, Campoalegre, and Algeciras [13] and in the department of Tolima in the municipalities of Chaparral and San Antonio [13,14], as well as in Abrego, Norte de Santander [15]. Additionally, *L. longiflocosa* is a species that presents highly anthropophilic and endophagic behaviors that incriminate it as a potential vector of *Leishmania sp* [16]. In this sense, this sandfly has shown a high susceptibility, under laboratory conditions, to infection with *Leishmania (V) braziliensis,* the main etiological agent of CL and its natural infection with *Le. (V) guyanensis* [17,18].

Considering the high relevance of these sandflies in the transmission of CL, it is important to highlight the scarce information on fundamental aspects of their biology, ecology, and bionomy [17,18]. Some relevant studies show the finding of individuals at immature stages (larvae) of *L*. *longipalpis* and *L*. *oswaldoi* in Brazil [19] and of *L*. *atroclavata* and *L. migonei,* mainly on the Colombian Caribbean coast, describing the abiotic characteristics of the probable sites of oviposition and breeding sites [20].

For most *Lutzomyia* species, the description and analysis of the parameters of spatio-temporal distribution, oviposition rates, hatching, survival-mortality, duration of immature stages, and other life cycle characteristics under uncontrolled conditions, are aspects not known in their entirety. The determination of these parameters in natural conditions will provide information that will allow estimates of their vector capacity and likely predict the times of greatest risk for contracting CL in endemic areas.

Previous studies show that for species of the verrucarum group, *L. spinicrassa, L. quiasitownsendi and L*. *youngi*, under standardized laboratory conditions, the average duration of their life cycle corresponds to 75.47; 69.85; and 61.07 days, respectively [9]. With respect to *L. longiflocosa*, classified within this group for its taxonomic characteristics, an average duration of 93.8 days for its life cycle under laboratory conditions has been reported [17], making this the longest cycle reported for the group of species studied.

Other studies have shown that, in natural circumstances in the sub-Andean zone of the departments of Tolima and Huila, *L. longiflocosa* presents a bimodal seasonal variation in the abundance of adults, showing two peaks: the first in the month of February and the second between the months of July and August. These patterns would be related to abiotic factors such as temperature, precipitation, and relative humidity, which fluctuate according to the climatic season [21, 22, 23] and, in turn, could have an influence on immature stages (eggs, larvae, and pupae), accelerating or slowing down their development (prolonged latency cycles) [22].

In order to provide knowledge about the biology of this species, taking into account its seasonality, the objective of this study was to establish the parameters of the life cycle of *L. longiflocosa* under semi-controlled conditions in a rural area of the municipality of Campoalegre, Huila during two seasons of the year.

## Methods

### Ethics statement

The people who captured the sandflies were not exposed to the bite of the insects, because they used protective clothing. Maintenance of *Lutzomyia* colonies including pre-reproductive and reproductive stages was approved by the ethics committee of the University of La Salle according to minutes of May 25, 2017. Field studies did not involve threatened or protected species.

### Study area

The study was carried out in the rural area of the municipality of Campoalegre, Huila, in the village "Venecia" located in mountainous territory on the western flank of the western mountain range, with coordinates 2°39′47"LN and 75°14′31"LW at an altitude of 1,618 m.a.s.l. (Fig 1). This area is characterized by an average temperature of 20°C and an annual rainfall of approximately 1,000 mm with a bimodal tendency, displaying two periods of high rainfall during the months of March and November [23].

**Fig 1.**
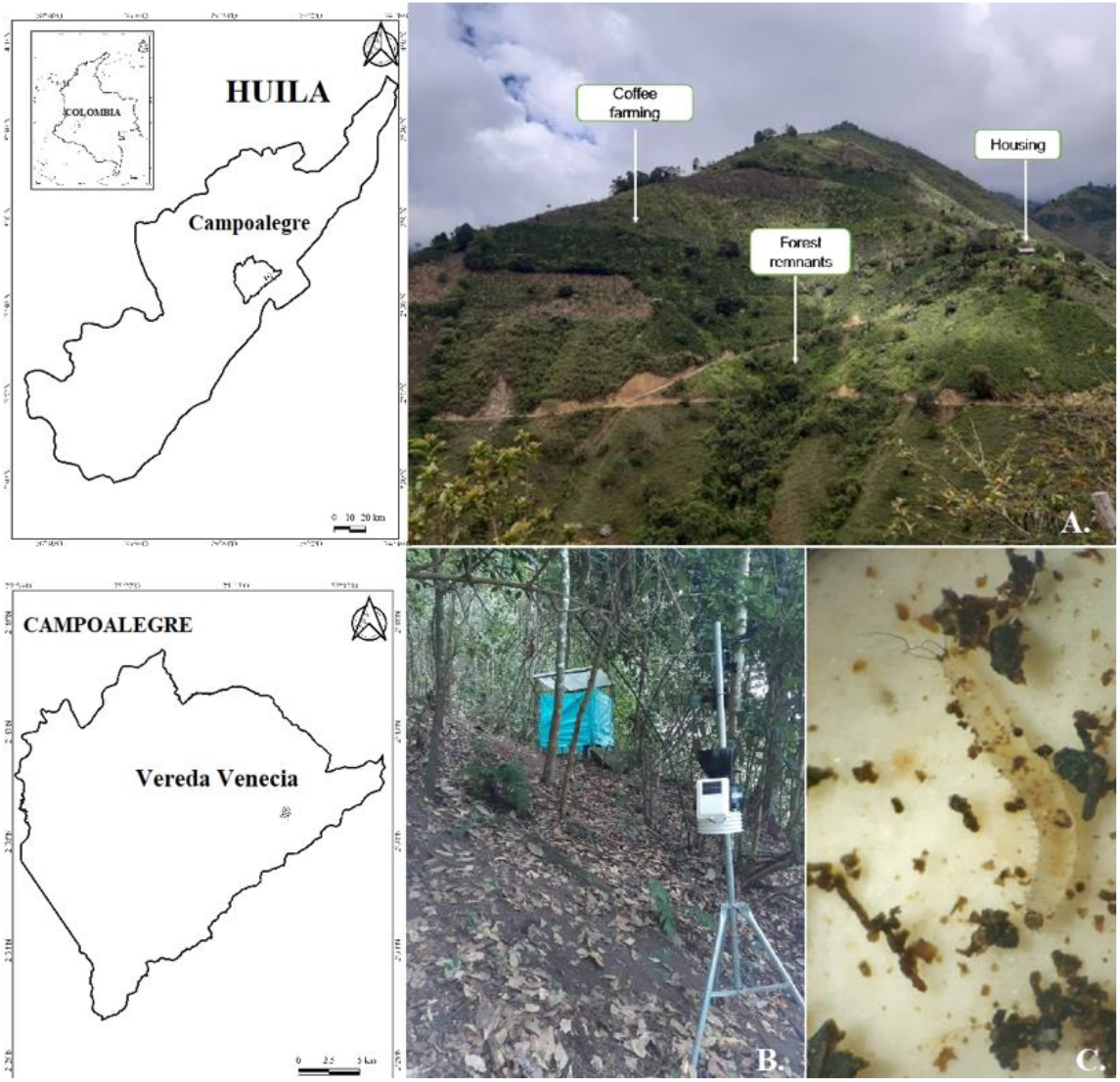
Geographical location of the rural area where the experiment was conducted. a) landscape of the area, b) installation of the monitoring area, c) microscopic view of stage II larvae (20x).

According to the Holdrige classification, this zone, corresponds to a premontane humid forest ecosystem [24]. The main economic activity of the area is associated with the use of land for the agricultural production of various products, especially for the cultivation of coffee (*Coffea arabica*).

### Sandfly collections

In order to include the climatic fluctuations that occur in the area during a year, two sandfly cohorts were monitored, each from the field collection of 100 female individuals of *L. longiflocosa*, in the months of February and July 2020, respectively. Cycle 1 (C1) was the follow-up carried out on the life cycle of the specimens collected in the first cohort between February and August 2020, and Cycle 2 (C2) was the follow-up carried out on the second cohort from July 2020 to January 2021.

The parental females were collected using protected human bait in forest relicts between 7:00 p.m. and 11:00 p.m., spaces that met previously reported criteria [22], such as a) presence of trees with a diameter at chest height > 30 cm, b) trees with rough bark, c) layer of undecomposed leaf litter > 5 cm, and d) previous determination of the presence of sandflies.

### Monitoring of life cycle parameters and developmental attributes

The insects collected were deposited into cages with metal frames (dimensions 30×30×30cm) and covered with cloth (muslin). The specimens were fed after their capture for 60 minutes, following the procedure of Mody and Tech [25] and Ferro et al. [26]. The fed females were individualized in transparent polyethylene brood cups (50 cm^3^ chambers) plaster-based to induce oviposition and sealed with a muslin shell. To enter and remove adult individuals, the muslin was centrally perforated with a hole 2 cm in diameter, sealed with a removable cotton plug. A water and 30% sugar solution were provided on the cover by means of two cotton swabs [27].

In the study area, the breeding sites were located within one of the previously chosen forests. The breeding chambers were arranged inside two styrofoam boxes, located 10 cm from the ground, for the subsequent beginning of the records of development in the conditions and factors of the habitat of this species (Fig 1). The variables humidity, temperature, and precipitation were recorded using a Davis Vantage Pro2 Weather Station® [28], located 4 m from the breeding site in the forest.

The larvae, in all their instar, were fed immediately after hatching up to (and including) instar IV through the regulated addition of food prepared from decomposing organic matter, following the protocol of Ferro et al. [26]. Emerging adults were individually placed in 25 cm^3^ brood cups, sugar solutions were provided, and longevity was registered.

For the identification of the specimens, the chemical clarification process described by Young in 1979 [27], and Maroli et al. in 1997 [29], was used to visualize internal keratinized structures (spermatheca and mature eggs). Subsequently, individuals were identified taxonomically following the taxonomic keys of Young and Duncan (1994) [30].

During the two cycles monitored in this study, the following parameters were registered: 1) Gonotrophic Cycle (GC), understood as the time elapsed from the blood intake of the female to the laying of the eggs, 2) Average duration in days of development of individuals per parental female at each stage of development, 3) Average of individuals per parental female at each stage of development, 4) Real fecundity (RF), expressed as the average number of eggs laid per parental female, 5) Potential Fecundity (PF), defined as RF plus the average number of eggs retained in the ovarioles-the determination of the number of eggs retained by each female was performed by dissection of the last segment of the abdomen and the subsequent counting of mature and chitinized eggs, 6) Fertility (F), defined as the percentage of eggs that hatched successfully, with respect to the total number of eggs laid per female [31], 7) Proportion of adults (males and females) emerged, and 8) Survival and mortality of individuals in the different immature stages (larvae I, II, III, IV, and pupa) and adults.

Similarly, attributes of population development were determined, defined as a) Hatching (I / H), I being the number of larvae I obtained, and H the number of oviposited eggs, b) Productivity (A / I), A being the number of adults obtained, c) Pupation (P / I), P being the number of pupae obtained, and d) Emergence of adults (A / P).

The duration of the different stages was determined by means of descriptive statistics, and the statistical differences between stages of development were evaluated with the Mann-Whitney Wilcoxon Test, using a P-value <0.05 as reference, with prior verification of the normality of the data. Finally, the correlations of Pearson and Spearman (according to the assumptions of normality) were considered to associate the life cycle variables obtained in the two development cycles studied with the averages of the abiotic variables recorded.

## Results

The life cycles monitored under semi-controlled conditions in the field showed significant differences in some of the parameters evaluated. C1 presented an average duration of 134.9 days, a shorter time (p<0.005) compared to C2, which displayed a duration of 148.78 days.

The GC in C1 presented a duration of 8.47 days on average, while in C2, it increased to 11.42 days (P=0.001). The development of immature stages, specifically in the larval stages, was more extensive for C2. This presented significant differences for larvae in stages II and III (P<0.001 and P = 0.008, respectively). The pupal stage in C1 presented a significantly longer time than C2 (P=0.003) (Table 1). Similarly, the longevity of mature individuals (adults) was more extensive during C2 (P=0.014).

**Table1.** Life cycle parameters and developmental attributes of *L. longiflocosa* in two seasonal life cycles under semi-controlled field conditions.

| Parameter | Life cycles monitored |  |  |
| --- | --- | --- | --- |
|  | C1 | C2 | P- value |
|  | Mean ± SD | Mean ± SD |  |
| Gonotrophic Cycle (GC) <sup>a</sup> | 8,47 ±3,18 | 11,42 ±2,21 | 0,001 <sup>a,c</sup> |
| Larval stage development (days) |  |  |  |
| Egg | 22,07±3,18 | 22,47 ±2.21 | >0,05 |
| Larvae I | 16,81±2,05 | 19,27 ±2,05 | >0,05 |
| Larvae II | 15,21±1,8 | 22,23 ±1,37 | <0,0001 <sup>c</sup> |
| Larvae III | 11,93±2,59 | 17,56 ±2,37 | 0,008 <sup>c</sup> |
| Larvae IV | 40,37±5,24 | 39,33 ±2,74 | >0,05 |
| Pupae | 24,48±3,93 | 22,9 ±1,03 | 0,001<sup>c</sup> |
| Female adults (<sup>b</sup>) | 4,1±1,88 | 6,94 ±0,93 | 0,001<sup>c</sup> |
| Male adults (<sup>b</sup>) | 4,24±2,3 | 6,03 ±1,03 |  |
| Average of individuals at each development stage |  |  |  |
| Egg | 15 ±2,73 | 12,64 ±1,87 | 0,021 <sup>c</sup> |
| Larvae I | 14,34 ±1,27 | 9,30 ±0,81 | 0,003 <sup>c</sup> |
| Larvae II | 13,55 ±1,15 | 9,17 ±0,37 | 0,008 <sup>c</sup> |
| Larvae III | 12,75 ±0,93 | 8,23 ±0,19 | 0,005 |
| Larvae IV | 11,82 ±0,51 | 7,93 ±0,15 | 0,008 <sup>c</sup> |
| Egg | 11,24 ±0,23 | 8,2 ±0,07 | 0,031 <sup>c</sup> |
| Female adults | 5,79 (±0) | 5,09 (±0) | >0,05 |
| Male adults | 5,44 (±0) | 4,78 (±0) |  |
| Fecundity y fertility |  |  |  |
| PF | 20 ± 1,43 | 26,03± 1,60 | 0,006 <sup>c</sup> |
| RF | 9,37 ± 1,15 | 5,02 ± 1,0 | 0,002<sup>c</sup> |
| F (%) | 47,52 ± 4,44 | 75,72 ± 18,40 | 0,016<sup>c</sup> |
| **Attributes of population development** |  |  |  |
| Hatching | 0,68 ± 0,37 | 0,76 ± 0,66 | 0.0006<sup>c</sup> |
| Productivity | 0,77 ± 0,17 | 0,57 ± 0,39 | 0.0009<sup>c</sup> |
| Pupation | 0,78 ± 0,17 | 0,57 ± 0,39 | 0.0009<sup>c</sup> |
| Emergence of adults | 1 | 1 | >0,05 |
| **Proportion of adults emerged** |  |  |  |
| Female (%) | 49,5 ± 19,60 | 65,074 ± 25,95 | 0,042<sup>c</sup> |
| Male (%) | 50,43 ± 19,60 | 34,92 ± 25,95 | 0,049 |

<sup>a</sup> Parental females.
<sup>b</sup> F1 adults fed sugar water only
<sup>c</sup> Statistically significant (P value < 0.005)

Regarding the number of individuals obtained in the F1 evaluated for C1 and C2, significant differences (P< 0.005), were found in the average number of individuals per parental female, specifically, during immature stages of development (larvae I, II, III, IV and pupae). When comparing the number of individuals per stage between the two cycles, it was observed that it was higher for C1 (Table 1), and significant differences were found in the stages of larval development: Larvae I (P = 0.003), Larvae II (P = 0.008), Larvae III (P = 0.005), Larvae IV (P = 0.008), and pupae (P = 0.031). However, no differences were observed in the final number of adult individuals (P>0.05) (Table 1).

In relation to reproductive indicators, the results in C2 show that F =75.72%/female, and PF =26.03 eggs/female, were higher than those found for C1 (F = 47.52%/female and PF= 20 eggs/female), with P=0.016 and P=0.006, respectively; however, RF showed an increase for C1 females (RF= 9.37/female), contrary to that presented for C2, (RF= 5.02/female) (Table 1).

Referring to attributes of development, the productivity, hatching, and pupation were higher in C1. The greatest difference was evidenced in the data obtained for hatching, in which it was 0.68 during C1 and for C2, it was 0.76 (Table 1). The emergence of the adult was the highest attribute in both cycles, presenting a higher value during C2.

Differences were observed in the survival presented between the two cycles monitored. For C1, all stages showed survival values greater than 70%, while for C2, the values were lower, starting with a survival percentage of 58% in the egg stage and increasing in each stage until reaching values greater than 90% from larvae III.

The mortality rate evidenced in C1 for the different stages was 32% (egg), 4% (larvae I), 6% (larvae II), 6% (larvae III), 7% (larvae IV) and 4% (pupae). For C2, the highest mortality rates occurred during the egg stages: egg, larvae I, and larvae II (24%, 41%, 27%, respectively) compared to later developmental stages like larvae III, larvae IV, and pupae (10%, 9%, and 3%).

On the other hand, the climatic variables recorded (precipitation, temperature, and relative humidity) fluctuated significantly during the development of the cycles evaluated. In the study area, the total rainfall was 1082 mm (min: 38.4mm, max: 164.6mm), evidencing a bimodal behavior defined by an increase during the month of March 2020 and those from November to January 2021. The average temperature was 18.76 °C (min:17.94 °C; max: 19.59°C); the increase in temperature coincided with the dry seasons presented in the month of February and those between August and October of the year 2020. Finally, the average relative humidity was 88.30% (min: 78.91%, max: 95.62%), which decreased considerably in the months of February, April, and those between August and October of the year 2020. Consequently, for C1, the number of average individuals per stage of development was positively correlated with temperature (R = 0.79, P = 0.024), and precipitation levels (R = 0.84, P = 0.012), while the average number of individuals of C2, presented a negative association with precipitation (R = −0.73, P = 0.033).

The larval development obtained in C1 did not present significant correlations with the environmental variables evaluated (temperature: R=-0.071, P>0.05, humidity: R=-0.32, P>0.05 and precipitation: R= −0.6, P>0.05); however, the survival rate of individuals revealed a negative association with the average precipitation levels (Fig 2).

**Fig 2.**
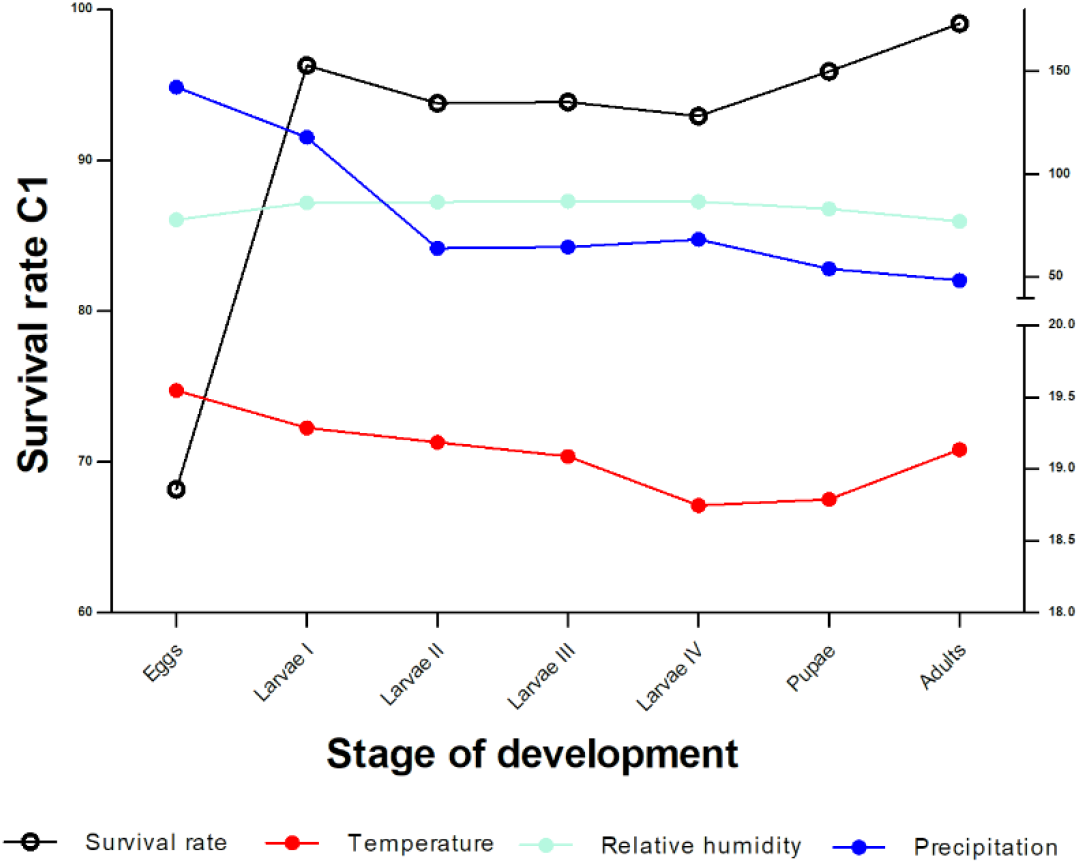
Survival rate of immature stages and related environmental variables for C1.

On the other hand, the larval development of C2 presented a positive and significant correlation between the survival rate and the average precipitation levels (R = 0.79, P = 0.03) and a probable association with humidity (R = 0.75, P = 0.05). Finally, it showed a negative but not significant association with temperature (R = −0.68, P>0.05) (Fig 3).

**Fig 3.**
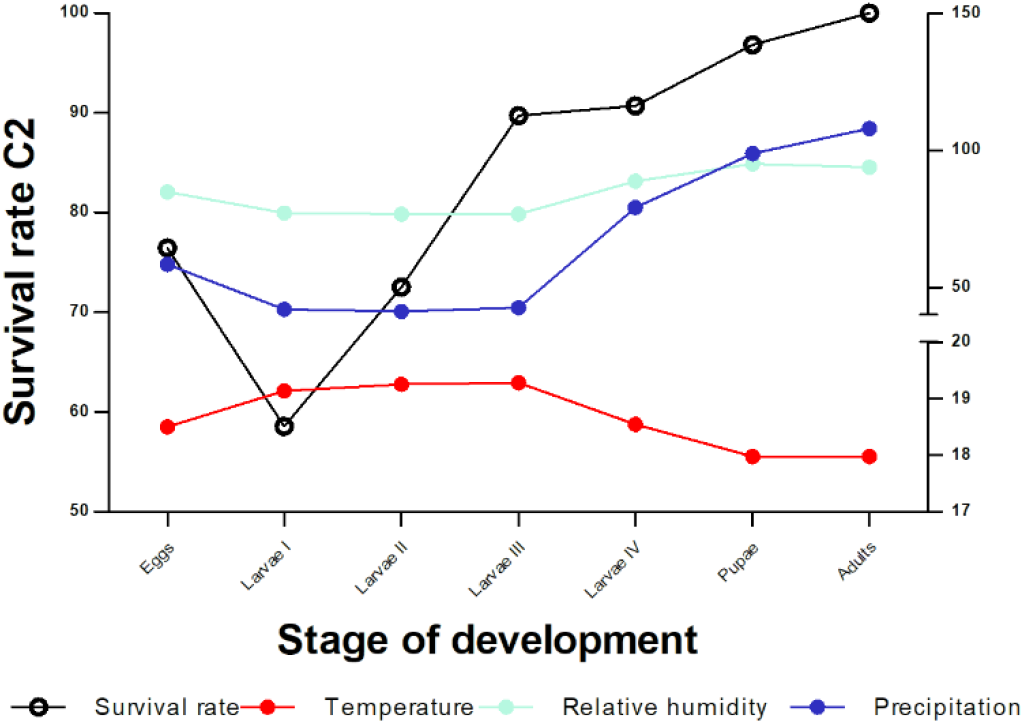
Survival rate of immature stages and related environmental variables for C2.

## Discussion

In this study, carried out in the rural area of Campoalegre, Huila, Colombia under semi-controlled conditions, it was possible to monitor the development of *L. longiflocosa* during two periods of the year, which together covered a full year, allowing to follow the development of two consecutive life cycles, starting from two cohorts of individuals of this species. Similarly, it was feasible to evaluate the incidence of precipitation, temperature, and relative humidity variables on the life cycle of *L. longiflocosa*.

Given the great difficulty of monitoring the development of these insects in their natural habitats, the possibility of evaluating how the conditions of the natural environment are determinants for the populations of this type of insects and, therefore, responsible for the prevalence of diseases such as leishmaniasis, is usually lost. In this work, although sandflies were protected in breeding chambers, environmental variables were not controlled. Consequently, the results obtained from life cycle parameters may represent an approximation to the conditions of development of *L. longiflocosa* in its natural habitat.

Since the first investigations in sandflies, the difficulty of detecting breeding sites has been evidenced due to several factors, namely a) the small size of the larvae, b) the great variety of possible sites for the laying of eggs and for the development of the different stages, and c) areas with high richness of decomposing organic matter, where there are diverse interactions of these insects with multiple tree species, generating countless microhabitats available for oviposition and development of pre-reproductive stages [20].

By virtue of the above, in recent decades, their colonization in controlled laboratory conditions has been proposed as the main strategy to clarify important aspects of their biology, life cycle, bionomy, genetics, physiology, and transmission of pathogens as well as for the evaluation of insecticides and new vector control methods [31,33]. Despite advances in sandfly colonization methods and strategies, only a little more than 20 of the approximately 1,000 species of sandflies reported have been colonized and kept by several generations in the laboratory. The main problems, associated with the difficulty of establishing colonies in the laboratory, are related to non-adaptation to laboratory conditions, which is reflected in the low reproduction rate of insects [33, 34], which makes this the main impediment to achieving a better knowledge of these species of interest in the aforementioned aspects. For this reason, studies related to the establishment and analysis of the life cycle of vector species of the genus *Lutzomyia* are scarce [18].

In particular, for *L. longiflcosa*, a single laboratory life cycle study conducted by Neira *et al*. in 1998 [17] is known, which shows the great difficulty of maintaining colonies of this species in the laboratory. In this study, they reported an average duration of the life cycle from egg to adult of 93.8 days, a value much lower than that obtained in this research. The differences found can be due to various causes, ranging from the type of substrate for growth of immature stages, the frequency and source of food, to the number of individuals per brood vessel, among other aspects [35]. However, given that environmental variables were not controlled in this study and insects were more exposed to environmental changes, it could be inferred that this exposure was the main cause of the differences found between the two studies. Similarly, Neira et al. [17] showed, as in the present research, that the longest development time was evidenced in larva stage IV. It is likely that the sandfly has the genetic capacity to extend this stage if the environmental conditions are not the most favorable because it must store enough energy to successfully develop the next stage (pupa), in which the individual does not feed and metamorphosis to adult. Similarly, it is plausible that some form of diapause has occurred, a phenomenon that has been reported in other *Lutzomyia* species. Diapause is considered an optional strategy expressed in response to environmental disturbances or scarcity of energy resources [35].

In the countries of Colombia, Brazil, and Venezuela, the life cycle under controlled laboratory conditions has been described for some species of the genus *Lutozmyia* involved in the transmission of the etiological agent of CL, including *L. spinicrassa, L. quasitownsendi, L. youngi* [9], *L. shanonni* [26], *L. evandroi*, *and L. longipalpis* [19], and *L*. *Ovallesi* [18]. Within the parameters estimated for these species, and according to the results obtained in this study, important differences were evidenced. *L. longiflocosa* displayed an average cycle length greater than 120 days compared to the 30-to-96-day fluctuation range established for the aforementioned species under controlled laboratory conditions. The hatching period of the eggs estimated in the previous studies takes place between 5 and 20 days, and the duration in the different larval stages occurs in periods between 15 and 70 days, obtaining the emergence of the adults between the following 7 to 14 days. These data contrast with those obtained in this study, in which *L. longiflocosa* presented a notable increase in the time required for each stage of development, taking up to 25 days for the hatching of eggs, 88 days in immature stages, and between 18 to 28 days for the emergence of adults after transformation into pupae (Table 1).

The reasons for these differences may be the same as those argued in this section in relation to the work of Neira et al. [17]; moreover, since these are species other than *L*. *longiflcosa* and, although they are phylogenetically close to each other, they may have enough genetic variations that cause different physical and behavioral responses to environmental changes.

In relation to the average GC obtained in this research, it fluctuated according to the cycle of the year evaluated (Table 1); however, it agrees with what was reported by Neira et al. [17] under laboratory conditions, with an average of 12.5 days. Studies related to GC are scarce, and even more so, when it is intended to determine the duration of the same in field conditions. Among the few studies, is relevant an investigation carried out in Brazil by Falcão et al. in 2017 [36], in relation to the duration of the GC of females of *Lutzomyia cruzi*. The procedure consisted of conducting two experiments with labeled, released, and subsequently recaptured females. In the first experiment of 172 released females, only two were recaptured five days after their release, without eggs or blood in the abdomen; while, in the second release, only one female was recaptured on the fourth day, which, two days later, laid eggs, suggesting a minimum GC of 6 days. The temperature during the experiments ranged between 17 and 33°C.

Regarding the production of individuals per female, Neira et al. [17] found that the average number of eggs obtained from *L. longiflocosa* under laboratory conditions was 27.6 eggs/female in contrast to the range obtained in this study with an average of 12 to 15 eggs/female (Table 1). It is likely that egg fertilization could be a determining factor in egg production probably due to early individualization and post-feeding confinement of females [26]. In 2021, Ribeiro Da Silva et al. [38] also reported that average egg production of individualized females was lower in the established colony of *L. longipalpis* than that of females who remained in groups (31.4 and 35.7, respectively).

In this study, fecundity showed values below 10, and in C2, the value was very low (5.02). These results are much lower than other studies that analyzed this parameter. In 2021, Ribeiro da Silva et al. [37] found an average fecundity of 33 in *L. longipalpis*; Escovar *et al.* [32] with *L. spinicrassa*, determined under laboratory conditions, a real fecundity of 36 and a potential fecundity of 37.4; Souza et al. in 2009 [34], with the complex of cryptic species of *L. longipalpis*, found fecundity values between 26.3 and 41.5, and Justiniano et al., in 2004 [38] with *L. umbratilis*, fecundity from 23.6 to 26.1.

Regarding fertility, in this research, important differences were evidenced between the two cycles monitored, finding the highest value in C2 with 75.72%, a value closer to that obtained by Neira et al. [17], who reported a fertility percentage of 82.7%, for *L. longiflocosa.* Other studies show even closer values of 75.5% [32] and 77% [38]. These differences in fecundity and fertility parameters between species may represent genetic differences in loci that control biological characteristics, given by geographical, reproductive isolation or mutational randomness [35].

The estimated developmental attributes for *L. longiflocosa* showed values higher than 0.5 in both cycles—0.77 for C1 and 0.57 for C2. Statistical analyses showed significant differences between the two cycles. Hatching was higher in C2, and pupation and productivity attributes (from adults) were higher in C1.

Some studies evaluating adult productivity of *Lutzomyia* in the laboratory show results with high variability. With *L. longipalpis*, in 2000, Luitgards-Moura et al. [39] found a minimum productivity of 21.1 in F1 and a maximum of 67.2 in F2. Ribeiro Da Silva et al. [37], determined productivity with values close to 25%. In this study, productivity was 57% in C2, and 77% in C1. These are high productivity values compared to the mentioned studies. It is likely that the differences presented are due to the fact that the uncontrolled conditions of this study allowed a better expression of the biological efficacy of these insects. Similarly, some of the disparities may be related to variations in the way this parameter is measured.

The lower survival rate of individuals for cycles C1 and C2 was evidenced during the immature stages, specifically in the egg stage, coinciding with what was evidenced for the species of the verrucarum group *L. spinicrassa, L. quiasitownsendi*, and *L. youngi*-[9]. On the other hand, Neira *et al*. [17] found in *L. longiflocosa* that the highest percentage of mortality occurred in larva IV, arguing that the natural extension of this stage increases the probability of fungal contamination and predation by mites. However, laboratory studies with other species of *Lutzomyia* show varied results in the stages with higher mortality; in larva I [32], egg and larva I [40], and egg and pupa [18]. In this sense, larval mortality is considered one of the main problems for the colonization of sandflies in the laboratory and, in particular, in the first stage, in which, due to their small size and low strength, individuals can be trapped in mycelia or in moisture condensation [33]. In contrast, Ribeiro Da Silva *et al.* [37], obtained a low mortality in pupae of *L. longipalpis*, arguing that the reason may be due to the fact that, at this stage, they do not feed and prefer dry environments in which invasion by fungi and other microorganisms is difficult.

The life cycles established for *L. longiflocosa* in the study area, covered seasonal periods associated with the fluctuation of precipitation in the Andean zone of Colombia. According to the correlations obtained, the survival of individuals could be positively associated with the increase in temperature and negatively with precipitation values (Fig 1 and 2). For the two cycles monitored in this study, the hatching of adults coincided with the low rainfall season (the month of August and those from December to February); and, as reported by Ferro et al [14], the number of adults of this species presents an increase in its abundance marked by the decrease in rainfall. Similarly, Rodríguez and Carvajal (2015) [11] also point out a negative association between the abundance of individuals and precipitation, based on a study carried out in 56 microhabitats located in the sub-Andean region of the upper Magdalena Valley between 100 and 2002 m.a.s.l., altitudinal conditions similar to those of this work.

On the other hand, for 5 years Valderrama et al. [41], evaluated different localities at medium elevations of the Colombian Andean region (1,000 to 2,000 m.a.s.l.) climatic variables as possible risk factors for the incidence of cutaneous leishmaniasis, associated with the presence of *Lutzomyia* species, concluding that the peak incidence of the disease is associated with a mean temperature of 20.6 °C (95% CI = 19.2–22.0 °C), a factor that, in turn, would be related to other variables such as vegetation cover and land use [11]. It should be noted that the average temperature recorded in this study was 18.76 ° C, very close to that reported in the aforementioned study, which allows us to infer a relationship between the abundance of *L. longiflocosa* and temperature. Consequently, the temporal distribution of this possible vector in areas of endemic transmission of CL would be conditioned by specific parameters of its life cycle in relation to the variation of the abiotic factors of their habitats.

In this study, significant differences were found between the cycles studied and in relation to several biological and reproductive aspects, demonstrating the effect of environmental variables on the development of the biological cycle of *L. longiflocosa* in its natural habitat. In this sense, the study of the relationships between environmental variables and reproductive parameters is essential to determine the entomological risk and the understanding of the transmission of leishmaniasis as well as to relate it to prevention activities.

The information obtained in this study allowed a greater depth of knowledge in the aspects of the population dynamics of *L. longiflocosa* in the area, which constitutes relevant information to optimize vector control measures.

## Supporting information

Dataset for table 1, figure 2 and figure 3.

## Acknowledgments

To Marco Fidel Suárez for his valuable management and dedication in the development of the field phase of this project.

## Notes

### Competing Interest Statement

The authors have declared no competing interest.

## References

1 Ovalle C, Londoño D, Salgado J, Gonzalez C. Evaluating the spatial distribution of Leishmania parasites in Colombia from clinical samples and human isolates (1999 to 2016). PLoS one. 2019; 14(3): 214–124. doi: 10.1371/journal.pone.0214124.

2 OPS. Leishmaniasis: Epidemiological Report of the Americas. No. 10, December 2021. Accessed February 7, 2023. Available at: https://iris.paho.org/handle/10665.2/53089.

3 Montoya LJ, Ferro C. Phlebotomes (Diptera: Psychodidae) of Colombia. In: Amat G, Andrade MG, Fernández F, editors. Insects of Colombia VII. Bogota: Colombian Academy of Exact, Physical and Natural Sciences. Jorge Alvarez Lleras Collection; 1999. P. 211-245.

4 NIH. National Institute of Health. Cutaneous, mucosal and visceral leishmaniasis event report, Colombia, 2018. Accessed February 7, 2022. Available at: https://www.ins.gov.co/buscadoreventos/Informesdeevento/LEISHMANIASIS_2018.pdf.

5 Velez I, Hendrickx E, Robledo S, Agudelo S. Gender and cutaneous leishmaniasis in Colombia. Cad. Saúde Pública. 2001; 17(1): 171–180. DOI: 10.1590/s0102-311X2001000100018.

6 Alvar J, Vélez I, Bern C, Herrero M, Desjeux P, Cano J. Leishmaniasis worldwide and global estimates of its incidence. PLoS One. 2012; 7(5): 124–135. https://doi.org/10.1371/journal.pone.0035671.

7 Lainson R, Rangel E. *Lutzomyia longipalpis* and the eco-epidemiology of American visceral leishmaniasis; with reference to Brazil - A Review. Memórias do Instituto Oswaldo Cruz. 2005; 100 (8): 811–827. https://doi.org/10.1590/S0074-02762005000800001

8 Pardo R, Ferro C, Lozano G, Lozano GA, Cabrera O, Davies C. Phlebotomine sandflies (Diptera: Psychodidae) vectors of cutaneous leishmaniasis and their ecological determinants in the coffee zone of the department of Huila XXVI Colombian Society of Entomology Congress. 1999. July 28-30, Bogotá.

9 Cabrera O, Ferro C. Life cycle of *Lutzomyia spinicrassa*, *L. quiasitownsendi* and *L. youngi*, species of the Verrucarum group (Diptera: Psychodidae). Actual Biol, 2000; 22(73): 225–232. https://doi.org/10.17533/udea.acbi.329660.

10 Osorno-Mesa E, Morales A, Osorno F, Muñoz P. Phlebotominae from Colombia (Diptera, Psychodidae). Description of *Lutzomyia longiflocosa* n.sp. and *Lutzomyia bifoliata* n. sp. Bol Mus Nat His Univ Fed Minas Gerais. 1970; (6). 18 p.

11 Rodríguez R, Carvajal A. Ecological Determinants of Forest to the Abundance of *Lutzomyia longiflocosa* in Tello, Colombia. Hindawi, 2015; (1):1–7. http://dx.doi.org/10.1155/2015/580718.

12 Santamaría E, Cabrera O, Avedaño J, Pardo R. Leg loss in *Lutzomyia longipalpis* (Diptera: Psychodidae) due to pyrethroid exposure: Toxic effect or defense by autotomy? J Vector Borne Dis. 2016; 53: 317–326 PMID: 28035108.

13 Pardo R, Cabrera O, Becerra J, Fuya P, Ferro C. *Lutzomyia longiflocosa*, possible vector in an area of cutaneous leishmaniasis in the sub-Andean region of the department of Tolima, Colombia, and the population’s knowledge about this insect. Biomedica. 2006; 26 (Suppl.1): 95–108.

14 Ferro C, Marin D, Gongora R, Carrasquilla M, Trujillo J, Rueda N, Valderrama A. Phlebotomine vector ecology in the domestic transmission of american cutaneous leishmaniasis in Chaparral, Colombia. Am J Trop Med Hyg. 2011; (85), 847–56. DOI:10.4269/ajtmh.2011.10-0560

15 Cárdenas R, Pabón E, Anaya H, Sandoval C. Presence of *Lutzomyia longiflocosa* (Diptera: Psychodidae) in the focus of tegumentary leishmaniasis in the municipality of Abrego, Norte de Santander. First record for the department. Clon, Institutional Journal of the Faculty of Health of the University of Pamplona. 2011; 3: 7–14.

16 Santamaria E, 2016. Effect of Industrially or Manually Treated Awnings with Long-Lasting Insecticides on the Vector Control of Cutaneous Leishmaniasis in the Sub-Andean Region of Colombia. Doctoral Thesis, Pontificia Universidad Javeriana, 2016. Available in: https://acortar.link/2fEKxR

17 Neira M, Diaz A, Bello F, Ferro, C. Laboratory study of the life cycles of *Lutzomyia torvida* and *Lutzomyia longiflocosa* (Diptera: Psychodidae) possible vectors of *Leishmania braziliensis* in the Colombian coffee region. Biomédica. 1998; 18(4), 251–265.

18 Cabrera O, Neira M, Bello F, Ferro C. Life cycle and colonization of *Lutzomyia ovallesi* (Diptera: Psychodidae), vector of *Leishmania* spp. in Latin America. Biomédica.1999; 19(3): 223–229. https://doi.org/10.7705/biomedica.v19i3.1026.

19 Freire de Melo M, Cunha J, Bezerra, S. Characteristics of the Biological Cycle of *Lutzomyia evandroi* (Costa Lima & Antunes, 1936) (Diptera: Psychodidae) under Experimental Conditions. Mem Inst Oswaldo Cruz. 2001; 96(6). DOI: 10.1590/s0074-02762001000600025

20 Vivero R, Torres C, Bejarano E, Cadena H, Estrada L, Florez F, Ortega E, Aparicio Y, Muskus C. Study on natural breeding sites of sand flies (Diptera: Phlebotominae) in areas of *Leishmania* transmission in Colombia. Parasites and vectors. 2015; 8 (116): 1–14. DOI: 10.1186/s13071-015-0711-y.

21 Morales D, Castano C, Lonzano E, Vallejo H. Description of the cutaneous leishmaniasis epidemic in Chaparral and San Antonio, 2003 and 2004 (week 24). Inf Quinc Epidemiol Nac. 9:180-6.

22 Carvajal L. Biotic and abiotic factors that partially define the abundance of *Lutzomyia longiflocosa*, vector of cutaneous leishmaniasis in the municipality of Tello, Huila. M.Sc. Thesis. Faculty of Sciences. Pontificia Universidad Javeriana.2008.

23 Ferro C, Lopez M, Fuya P, Lugo L, Cordovez J, Gonzalez C. Spatial Distribution of Sand Fly Vectors and Eco-Epidemiology of Cutaneous Leishmaniasis Transmission in Colombia. PLoS One. 2015; 10(10): 1–16: https://doi.org/10.1371/journal.pone.0139391

24 Holdridge L. Life Zone Ecology. Tropical Science Center. San José, Costa Rica. 1a. ed: IICA, 1982.

25 Modi G, Tesh R. A simple technique rearing *Lutzomyia longipalpis* and *Phlebotomus papatasi* (Diptera: Psyhodidae) in the laboratory. Journal of Medical Entomology. 1983; 20; 568–69. DOI: 10.1093/jmedent/20.5.568

26 Ferro C, Cardenas E, Corredor D, Morales A, Munstermann L. Life Cycle and Fecundity Analysis of *Lutzomyia shannoni* (Dyar) (Diptera: Psychodidae). Mem. Inst. Oswaldo Cruz. 1998; 93(2): 195–200. https://doi.org/10.1590/S0074-02761998000200011.

27 Young D, Morales A, Kreutzer R, Alexander B, Corredor A, Tesh R. Isolation of *Leishmania braziliensis* from cryopreserved Colombian sandflies (Diptera: Psychodidae). J Med Entomol. 1987; 24:587. https://doi.org/10.1093/jmedent/24.5.587

28 Davis instruments. Software WheatherLink Vantage Pro2. Windows. Fecha de consulta: 10 de febrero del 2020. Disponible en: https://www.weatherlink.com/.

29 Maroli M, Feliciangeli M, Arias J. Methods of capture, preservation, and collection of sandflies (Diptera: Psychodidae). Pan American Health Organization (OPH); 1997.

30 Killick-Kendrick R. Guide to the identification and geographic distribution of *Lutzomyia* sand flies in Mexico, the West Indies, Central and South America (Diptera: Psychodidae). Trans R Soc Trop Med Hyg [Internet]. 1995;89(1):125. http://dx.doi.org/10.1016/0035-9203(95)90687-8.

31 Rabinovich JE. Ecology of Animal Populations. Regional Program for Scientific and Technological Development; 1978.

32 Escovar, J., Bello, F., Morales, A., Moncada L., Cardenas, E. Life Tables and Reproductive Parameters of *Lutzomyia spinicrassa* (Diptera: Psychodidae) under Laboratory Conditions. Mem Inst Oswaldo Cruz. 2004; 99 (6). DOI: 10.1590/s0074-02762004000600012

33 Lawyer P., Killick-Kendrick M., Rowland T., Rowton E., Volf P. Laboratory colonization and mass rearing of phlebotomine sand flies (Diptera, Psychodidae). Parasite. 2017: 24 (42). http://dx.doi.org/10.1051/parasite/2017041.

34 Souza, N. A., Andrade-Coelho, C. A., Silva, V. C., Ward, R. D., & Peixoto, A. A. Life cycle differences among Brazilian sandflies of the *Lutzomyia longipalpis* sibling species complex. Medical and Veterinary Entomology. 2009; 23(3), 287–292. https://doi.org/10.1111/j.1365-2915.2009.00818.x.

35 Mann, R. S., & Kaufman, P. E. Colonization of *Lutzomyia shannoni* (Diptera: Psychodidae) utilizing an artificial blood feeding technique. Journal of Vector Ecology. Journal of the Society for Vector Ecology. 2010: 35(2), 286–294. https://doi.org/10.1111/j.1948-7134.2010.00084.x.

36 Falcão de Oliveira, E., Oshiro, E. T., Fernandes, W. de S., Murat, P. G., de Medeiros, M. J., Souza, A. I., de Oliveira, A. G., & Galati, E. A. B. Experimental infection and transmission of *Leishmania* by *Lutzomyia cruzi* (Diptera: Psychodidae): Aspects of the ecology of parasite-vector interactions. PLoS Neglected Tropical Diseases. 2017: 11(2), e0005401. Doi: https://doi.org/10.1371/journal.pntd.0005401.

37 Ribeiro da Silva, R. C., Nava Piorsky Dominici Cruz, L., da Silva Coutinho, J. M., Correia Santana, N. C., & Macário Rebêlo, J. M. Maintenance and productivity of a Lutzomyia longipalpis (Diptera: Psychodidae) colony from an area endemic for visceral and cutaneous leishmaniasis in Northeastern Brazil. Journal of Medical Entomology. 2021: 58(4), 1917–1925. https://doi.org/10.1093/jme/tjab053.

38 Justiniano, S. C. B., Chagas, A. C., Pessoa, F. A. C., & Queiroz, R. G. Comparative biology of two populations of *Lutzomyia umbratilis* (Diptera: Psychodidae) of Central Amazonia, Brazil, under laboratory conditions. Brazilian Journal of Biology. 2004: 64(2), 227–235. https://doi.org/10.1590/s1519-69842004000200007.

39 Luitgards-Moura, JF, Castellón Bermúdez, EG, & Rosa-Freitas, MG. A Aspects Related to Productivity for Four Generations of a *Lutzomyia longipalpis* Laboratory Colony. Memorias del Instituto Oswaldo Cruz. 2000: 95 (2), 251–257. https://doi.org/10.1590/s0074-02762000000200021.

40 Castillo, A., Serrano, A. K., Mikery, O. F., & Pérez, J. Life history of the sand fly vector *Lutzomyia cruciata* in laboratory conditions. Medical and Veterinary Entomology. 2015: 29(4), 393–402. https://doi.org/10.1111/mve.12127.

41 Valderrama-Ardila, C., Alexander, N., Ferro, C., Cadena, H., Marín, D., Holford, T. R., Munstermann, L. E., & Ocampo, C. B. Environmental risk factors for the incidence of American cutaneous leishmaniasis in a sub-Andean zone of Colombia (Chaparral, Tolima). The American Journal of Tropical Medicine and Hygiene. 2010: 82(2), 243–250. https://doi.org/10.4269/ajtmh.2010.09-0218.

